# Herpetologists’ Conservation Research Focus Drives Their Intentions to Participate in Future Public Engagement

**DOI:** 10.1101/2021.05.14.444251

**Authors:** Kirsten A Hecht, Kathryn A. Stofer, Martha Monroe, Geraldine Klarenberg, Max A. Nickerson

## Abstract

Public Engagement with Science (PES) is a popular topic in the science community due to general concerns about public support for science, attitudes toward science, and changes in scientific funding requirements. PES may be especially relevant in conservation disciplines as the public plays an important role in conservation practice. Herpetofauna specifically stand to benefit, as PES activities can help improve attitudes and conservation behavior of participants toward uncharismatic species. We assessed the current scope of herpetologists’ PES activities and investigated factors associated with their participation in PES. We used a closed-ended question survey distributed via the listservs of four American herpetological organizations. Herpetologists’ intentions to engage at least 10 hours in the next 12 months significantly differed between herpetologists with high and low conservation research focuses, but hours of engagement in the past 12 months was not significantly different among these groups. Despite most responding herpetologists having limited formal training, time, resources, and institutional support, many participated in a variety of PES activities, often utilizing partnerships and their own resources. Sampled herpetologists rarely evaluated their PES activities or considered publishing about their engagement activities. Some respondents expressed unease with the idea of message framing. Respondents were interested in evaluation training and providing accessible opportunities, and grant funds were the most likely interventions to increase herpetologists’ participation in PES. These results provide reference data and insight into the public engagement practices and needs of practicing herpetologists and conservation scientists.

**Author statement:** We informed participants of their rights and protections as approved by University of Florida IRB201800258. None of the authors have conflicts of interest related to this research. Funding for the research was provided by a Roger Conant Grant from the Society for the Study of Amphibians and Reptiles (SSAR). SSAR had no role in interpretation of data, the writing of the report, or the decision to submit the article for publication.

## Introduction

As a field, conservation biology is intimately tied with public interest. Wildlife conservation efforts based on biological scientific knowledge and solutions alone are often unsuccessful at meeting conservation goals (Agrawal and Gibson, 1999; Knight et al., 2006; Mascia et al., 2003; Meffe, 2002; Wilson et al., 2007) and there have been repeated recent calls to not only include, but to make social factors a key aspect of conservation (Ban et al., 2013; Bennett et al., 2017; O’Donnell & Durso, 2014; Saunders et al., 2006; Wake, 2008b). Mascia et al. (2003, p. 649) expressed the essence of this shift in the following passage, “Although it may seem counterintuitive that the foremost influences on the success of environmental policy could be social, conservation interventions are the product of human decision-making processes and require changes in human behavior to succeed.” Public engagement with science (PES), which the American Association for the Advancement of Science (AAAS) describes as “intentional, meaningful interactions that provide opportunities for mutual learning between scientists and members of the public,” (AAAS, 2021; para. 1) is one tool that could bridge between these ecological and socio-ecological factors, potentially serving to increase public support for policy and conservation activities.

A holistic approach to conservation may be particularly important to protecting less charismatic animal taxa that receive less support from both the public and from the wildlife field. Reptiles and amphibians in particular face unique conservation challenges when compared to other organismal groups. Herpetofauna continue to decline in number with researchers now estimating that over 40% of the world’s studied amphibians (IUCN, 2018) and ~20% of studied reptiles (Böhm et al., 2013) are threatened with extinction worldwide while also facing localized decreases in occupancy and metapopulations (Adams et al., 2013; Grant et al., 2016). Herpetofaunal conservation also faces challenges from deep social and cultural barriers including attitudes, social values, and norms (Alves et al., 2012; Ceríaco, 2012; Ceriaco et al., 2011; Perry-Hill et al., 2014; Tarrant et al., 2016), negative emotions such as disgust (de Pinho et al., 2014; Gunnthorsdottir, 2001; Knight, 2008), and even innate fear (Hoehl et al., 2017; LoBue and DeLoache, 2008), which ultimately lead to high rates of persecution, less public support for conservation, and less priority in regards to research and conservation attention and funding (Bonnet et al., 2002; Clucas et al., 2008; Gratwicke et al., 2012).

Public engagement activities provide an opportunity for herpetologists to increase conservation support for reptiles and amphibians among the public. Interventions focused on general knowledge (i.e., one-way communication) or simple exposure, such as watching snakes in a zoo, have not shown significant changes in attitudes towards herpetofauna (Prokop and Tunnicliffe, 2010; Tomažic, 2011), or changes in values or behavior (Morgan and Gramann,1989). In contrast, positive interactive experiences with snakes, specifically direct contact (Prokop and Tunnicliffe, 2010) and modeling, where individuals can see others positively interacting with animals (Morgan and Gramann, 1989), were most effective at eliciting changes in participants, particularly when information was included as part of the experience. Even limited one-day exposures, such as field trips where students were able to handle and measure animals, altered likeability and individual support for conservation of snakes and salamanders (Morgan & Gramann, 1989; Reynolds et al., 2018). Programs utilizing a full treatment set (exposure, knowledge, modeling, and direct contact) may have the highest success. A citizen science program in India, for example, reduced human persecution of snakes in the area and improved attitudes of participants towards snakes (Balakrishnan, 2010). Ultimately these types of activities may help change attitudes about uncharismatic species because they foster familiarity with the groups or species and can even change aesthetic beliefs and attitudes which has been linked to increased positive attitudes towards species (Jimenez and Lindemann-Matthies, 2015; Reimer et al., 2014).

Research related to PES suggests that in some cases, a scientists’ field or topic of study may be an important factor in determining whether they choose to engage with public. For example, a Pew Research Report found that scientists who felt the public had more interest in their research area were more likely to engage (Pew Research Center, 2015). In that same study, scientists who felt there was controversy in their fields were more likely to talk to reporters and citizens as well as use blogs and social media when compared to scientists who rarely or never saw debate about their field in the public eye (Pew Research Center, 2015). In another study of 1,254 biomedical scientists, Dudo (2012) found that respondents who believed that publicly communicating about their work was beneficial to society were more likely to engage in public activities. However, it is unclear whether conservation sub-fields follow this trend. Despite the potential value of public engagement as a conservation tool, there remains limited study on the broader public engagement activities of conservation biologists focused on reptiles and amphibians. During a literature review, we found only limited examples of publications related to education and outreach in any U.S. based herpetological journal over the past 20 years (Clancy et al., 2021). These publications largely focused on herpetology courses in formal education settings or museums (Chiszar, 1998; Frost, 1998) and citizen science from a data collection perspective (O’Donnell and Durso, 2014; Weber et al., 2016). A handful described education and conservation outcomes for a variety of audiences including herpetological educators, K- 12 teachers, and children (Ballouard et al., 2012; Gangloff, 2011; Rommel-Crump et al., 2016; Wojnowski, 2008). A non-peer reviewed report found surveyed individuals in all sectors of herpetology, including hobbyists, participating in public outreach activities, specifically helping run or organize public education programs alone or on behalf of herpetological societies (Southwestern Center for Herpetological Research, 2017). Many of the surveyed recreational and professional herp-enthusiasts have attended or facilitated some type of educational activity. There is no additional research to our knowledge specifically related to public engagement by herpetologists, including whether herpetologists whose research focuses primarily on conservation are more likely to engage with the public due to a higher public interest factor. Therefore, we surveyed herpetologists regarding their public engagement activities and experiences as a case study to help improve the effectiveness of public engagement as a tool for the conservation of uncharismatic animals. The objectives of this study were to determine:

1. The types of engagement activities in which herpetologists participate;
2. How prepared herpetologists are to engage with the public;
3. The barriers herpetologists face to engage with the public; and
4. Whether herpetologists’ professional interest in conservation is a significant predictor of their public engagement participation.

## Methods

### Population of Interest

The population of interest for this study is herpetological researchers, ages 18 and over, who reside in the United States. We defined herpetological researchers as individuals who conduct original scientific herpetological research and have published at least one related peer-reviewed paper in a scientific journal. The population encompassed herpetologists from all potential sectors, including, but not limited to, universities, zoos, government agencies, museums, non-governmental organizations, and private industry, and contained herpetologists with varied levels of herpetological related conservation research.

### Survey Instrument

We developed a survey consisting of 37 closed-ended questions which was based on our research questions as well as results from fifteen semi-structured interviews (Hecht, 2021). Questions covered demographics; herpetology research interests; how often conservation was a focus of their work; study animals; and public engagement knowledge, training, and practices. The survey had four screening questions that asked if respondents fit the criteria of our population of interest (residing in United States, 18 or older, conducted original research on reptiles and/or amphibians, and have published at least one peer-reviewed paper on reptiles and amphibians) and a general question about what types of publicly oriented activities they had participated in within the last 12 months. Following these questions, we provided the AAAS definition of public engagement with science as “intentional, meaningful interactions that provide opportunities for mutual learning between scientists and members of the public.” (AAAS, 2021; para. 1) We measured level of participation in public engagement by asking about the frequency of their participation in the last 12 months as well as their intention to participate in the next 12 months. To quantify past engagement, participants were asked to choose categories corresponding to how many hours of herpetology related public engagement they participated in as an expert within the last 12 months (I did not engage, <10, 10-49, 50-99, and >100). Future participation was measured from two 7- point bipolar adjective scale questions (Ajzen, 2006) which measured their intention to participate in at least 10 hours of public engagement over the next 12 months: 1) I intend to participate in public engagement as a herpetology expert for at least 10 hours in the next 12 months (highly likely to highly unlikely), and 2) I will try to participate in public engagement as a herpetology expert for at least 10 hours in the next 12 months (definitely will not to definitely will).

Eight graduate students and three faculty with expertise in questionnaire design, public engagement, and/or herpetology fields reviewed the survey in two rounds. We then pilot tested the survey with two students and four professionals affiliated with environmentally related fields that would be familiar with the terminology, but not within our specific population of interest, i.e., non-herpetologists, to maximize the number of people we sampled, as most herpetologists would receive the survey through their professional organizations. After each round, participants sent us either verbal or written feedback, and we incorporated their feedback into the survey. The survey was entered into Qualtrics prior to distribution.

### Recruitment

We sent the final Qualtrics survey link to members of four North American herpetological societies and organizations to share via an anonymous link sent through their membership listservs and newsletters: Partners in Amphibian and Reptile Conservation (PARC) (membership unknown), the Society for the Study of Amphibians and Reptiles (SSAR) (1,613 members), Herpetologists’ League (HL) (599 members in 2015), and the American Society for Ichthyologists and Herpetologists (ASIH) (1,517 members with approximately half identifying as herpetologists). The survey was open for one month between 1 September, 2019 and 1 October, 2019. Every 25th participant was eligible to receive a $10 prepaid debit card by providing their email in a separate survey provided upon the competition of their survey.

### Analysis

Prior to analysis, we converted the 7-point bipolar scales of the intention questions to integers ranging from −3 to 3 and added the two intention questions together to obtain a composite score for each participant. We checked all variables for normality and collinearity prior to analysis. A correlation of greater than 0.6 between predictor variables was used as the cut off value for determining multicollinearity (Dormann et al., 2013). We used basic descriptive statistics to analyze survey data for all objectives. To determine if herpetologists working in the conservation biology subfield were more likely to engage with the public, we analyzed subfield data and conservation focus of research data using ordinal regression analysis. We used a backward stepwise model building approach using Akaike Information Criterion (AIC) estimates to determine the best fit for each individual model (Akaike, 1998). We also examined whether the level of conservation focus of respondents impacted their engagement activities. Due to the low response count in the never category (n=8), responses to the question regarding the amount of research related to herpetological conservation was transformed from 5 choices (Never (n=8), Less than half the time (n=47), About half the time (n=23), More than half the time (n=45), and Always (n=55) to 3 categories (Low (n=55), Medium (n=23), and High (n=100)) prior to analysis. We hypothesized that individuals who had a high conservation focus in their research would a) have participated more hours in public engagement activities in past 12 months and b) would also exhibit a higher intention to participate in public engagement for at least 10 hours in the next 12 months. Due to the lack of normality in the data, we conducted a non-parametric Kruskal-Wallis test to look for differences among the three conservation focus categories to test our hypotheses. We then conducted a post-hoc pairwise comparison using a Dunn test with a Bonferroni correction. All significance tests used an alpha value of 0.05.

We analyzed all data using R (R Core Team 2020). We conducted ordinal regression analysis using the polr and StepAIC functions in the MASS R package (Venables & Ripley, 2002); We used the FSA R package (Ogle et al., 2020) to conduct the Dunn test. All other stats were analyzed through the base stats program of R.

## Results

A total of 355 unique individuals started the survey, of which 217 were eligible participants that passed the four screening questions. Of those respondents, 178 completed the full survey. Due to disseminating the survey through professional associations and overlapping memberships, it was not possible to calculate a population size or response rate. Estimated survey time from Qualtrics was 17.4 minutes and pilot testers took between 14 and 43 minutes. Only 12 study participants took longer than 1.5 hours to complete the survey. The mean survey duration for the remaining 166 participants was 21.15 ± 13.00 minutes.

### Demographics

Demographics matched findings of other demographic surveys of herpetology organizations (*2019 Diversity Survey Report*, 2020; *Demographic and Atmospheric Survey*, 2020). Participants included representatives from all adult age categories with 57.9% coming from two age classes (25-34 and 35-44 years; Figure 1). Most survey takers self-identified as men (58.4%), with the remaining participants identifying as women (38.8%) and non-binary/third gender (1.1%). A small number (1.7%) preferred not to disclose their gender. Of the participants, 7.3% identified as Hispanic, Latino, or Spanish Origin. Most participants (89.9%) described their race as white, while 1.7% and 0.5% identified as Asian and Black/African American, respectively. Three respondents described themselves as multi-racial.

**Figure 1.**
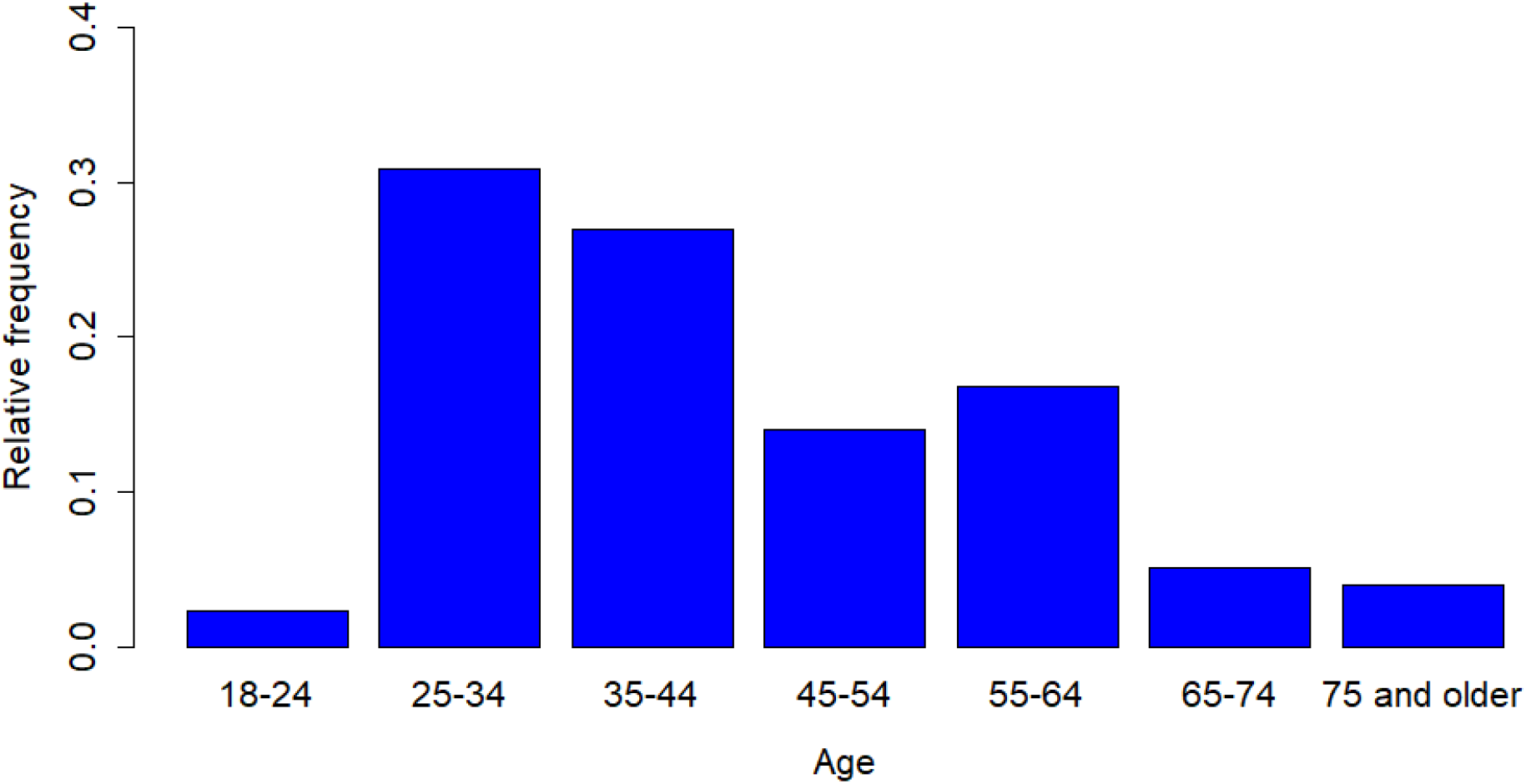
Age distribution of respondent herpetologists.

Respondents ranged across experience levels, but 41.0% had been in the field for at least 20 years. Most participants had an advanced degree with 55.6% holding some type of doctoral degree, 34.3% holding a master’s degree, 8.4% having a bachelor’s degree, and 1.7% having a high school diploma as their highest education level. Participants had expertise in a variety of sub-fields, with many selecting multiple sub-fields to describe their research. The number of identified sub-fields per participant ranged from 1 to 8, with a mean 3.32 ± 1.41 sub-fields. Ecology and Conservation were the most frequently selected categories with over 70% of participants identifying either of them as a sub-field of their research. Behavior (42%) and Evolution (40%) were the next most frequently reported sub-fields. Participants also represented a variety of sectors, with almost a third coming from public institutions. Individuals from undergraduate- and graduate-serving institutions represented the highest percentage of individuals, over 30% of the sample participants, followed by agencies (22%), and museums (10%). One out of five respondents reported themselves as students. Survey takers represented all four of the organizations surveyed: ASIH (54.5%), HL (44.4%), SSAR (70.2%), and PARC (49.4%). Many individuals were members of multiple societies, with respondents belonging to a mean of 2.19 ± 1.10 societies and 11.8% of respondents belonging to all four. In addition, 27.0% of responding herpetologists belonged to a regional herpetological society and 11.2% identified as members of a local herpetological organization.

### Objective 1: Types of Activities

Almost all (98%) of those surveyed had participated in public engagement activities as a herpetology expert, and 36% of respondents reported public engagement was part of their job duties. Within the last 12 months, 67.5% of participants reported they had participated in at least 10 hours of public engagement activities, with 52.2% citing 10-49 hours and 10.2% reporting 55-99 hours of public engagement activities in the same period. Only 3.6% of respondents did not participate in any public engagement activities as a herpetology expert in the last 12 months. Half of the participants also had high levels of intention to participate in at least 10 hours of public engagement in the next 12 months with 47.8% expressing they were both highly likely to participate and also intended to participate at this level.

Respondents were more likely to develop their own engagement programs at least sometimes (85.2%) rather than participate in an already established program (62.5%). All but 6.5% of participants parterned in at least one stage of engagement, with 37% partnering during activity development and 48% for implementation. Only 14% of respondents never use live animals in their engagement activities, while 35.2% and 18.2% use them sometimes and always, respectively. The number of public-oriented activity types that respondents were involved with over the past 12 months ranged from zero to fifteen with a mean of 4.65 ± 3.06. In-person lectures for the public, social media, and citizen science were the three most common public-oriented activities (Figure 2). Virtual lectures, blogs, science cafés, and teacher trainings were the least reported activities.

**Figure 2.**
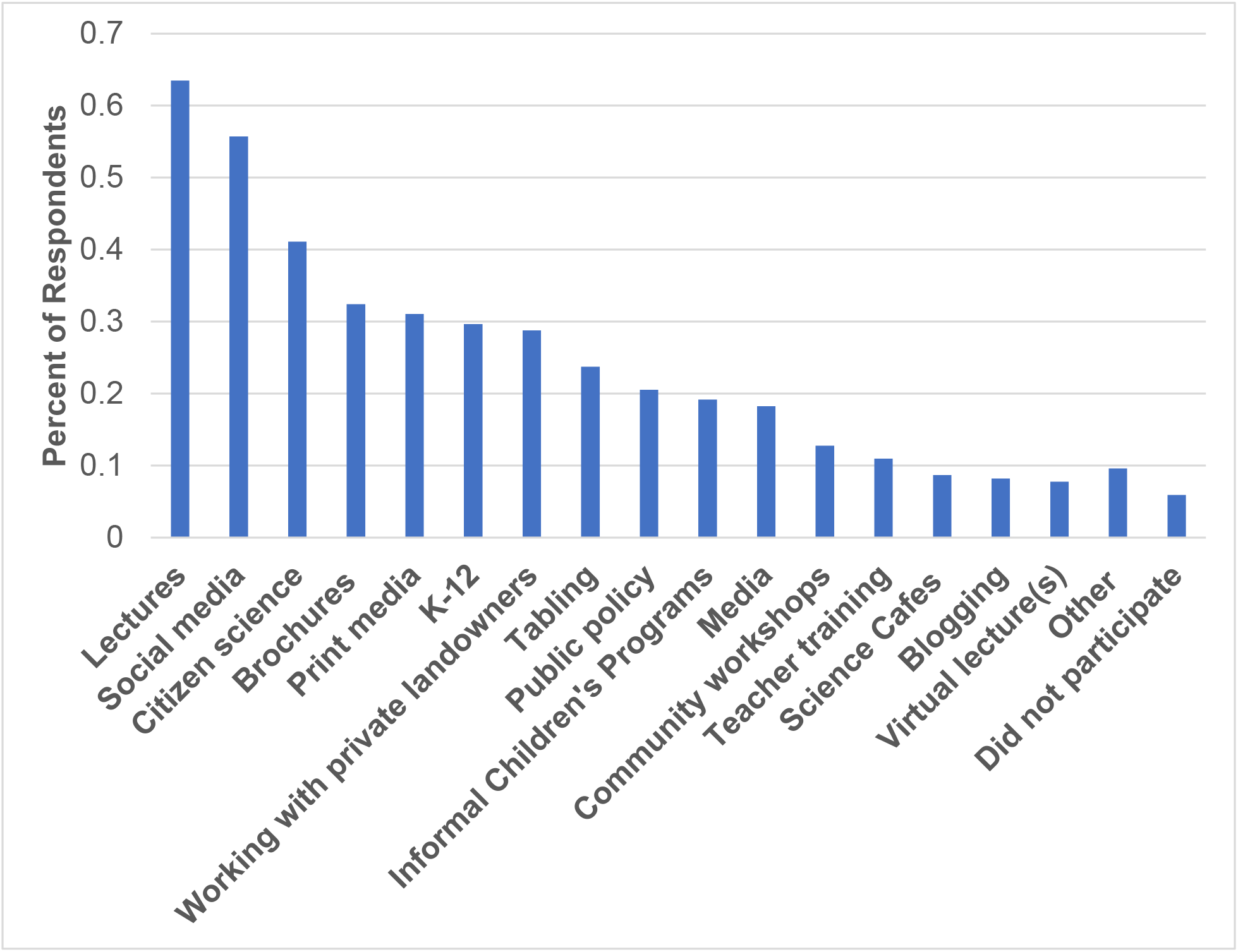
Types of public activities undertaken by respondent herpetologists within the past 12 months.

### Objective 2: Training and Knowledge

Approximately two out of three participants (67.6%) stated they had never had any formal training in public engagement. Similarly, few respondents regularly used any resources to increase their knowledge about engagement, with 71.54% and 52.3% rarely or never reading peer-reviewed literature or using other resources to improve their public engagement strategies, respectively. Most participants reported that they adjust their message based on the audience they were engaging with, with 46% doing this all the time compared to only 4% never doing so. Three out of four respondents reported that they rarely or never evaluate their public engagement activities, and 54% have never thought about publishing about their public engagement activities. For those that do evaluate, evaluation was largely limited to less systematic methods like observing participants (43.3%), participant comments (46.1%), or counting the number of participants (34.8%). Respondents rarely evaluate programs with opinion surveys (16.9%), knowledge tests (7.9%), interviews (5.1%), social media metrics (11.8%), and/or external evaluators (8.4%). However, evaluation training was the most frequently selected option for what type of training herpetologists would be interested in (44.4%). How to have difficult conversations (38.8%) and how to develop programming (36.0%) were the next two most popular training choices. About one in five (21.9%) of respondents were not interested in additional training.

### Objective 3: Barriers

Participants cited limited time (74.3%) as the most common factor that at least sometimes prevented their participation in public engagement activities, followed by funds (46.9%). On a 7-point rising scale, 62.0%, 64.5%, and 62.5% of respondents reported that training, skills, and work very rarely or never prevented their participation respectively. Most participants used their own resources more often than work resources during their public engagement activities. When asked what would make them most likely to engage more often, grants or funds for public engagement were cited the most often (25.8%) followed by more engagement opportunities (23.0%) and dedicated work time (22.5%). Work-related recognition (9.6%), potential for publications (4.5%), and training (2.2%) were not commonly cited as factors that would make participants engage with the public more often.

### Objective 4: Impact of Conservation

Respondents varied in how much they focus on conservation in their herpetology research, with 30.9% citing conservation as a focus all the time and 26.4% reporting they do conservation-related research less than half the time. Most respondents (69.1%) focus on herpetofauna conservation research at least half the time. Using a Kruskal-Wallis test, we found that individuals from these three levels of conservation research (low, medium, high) did not differ significantly in the hours of engagement in the past 12 months (**χ**^2^=4.92, df=2, p-value=0.086). In contrast, intention to engage did differ significantly among the three conservation levels (**χ**^2^=7.38, df=2, p-value=0.025, Figure 3). In follow-up pair-wise comparisons using a Dunn’s Test, we found a significant difference between the low and high conservation groups regarding intention to participate in at least 10 hours of public engagement in the next 12 months (Z=2.42, p-value= 0.049). Following Bonferroni correction, we did not find a significant difference between medium and high levels of conservation focus (Z= 2.0433, p-value= 0.123). We found that sub-fields were not significant predictors of future intentions to engage at least 10 hours over the next 12 months, but conservation biology, taxonomy, behavior, and education were significant terms in the ordinal logistic regression predicting how many hours over the past 12 months respondents engaged with the public (Table 1; Table 2).

**Figure 3.**
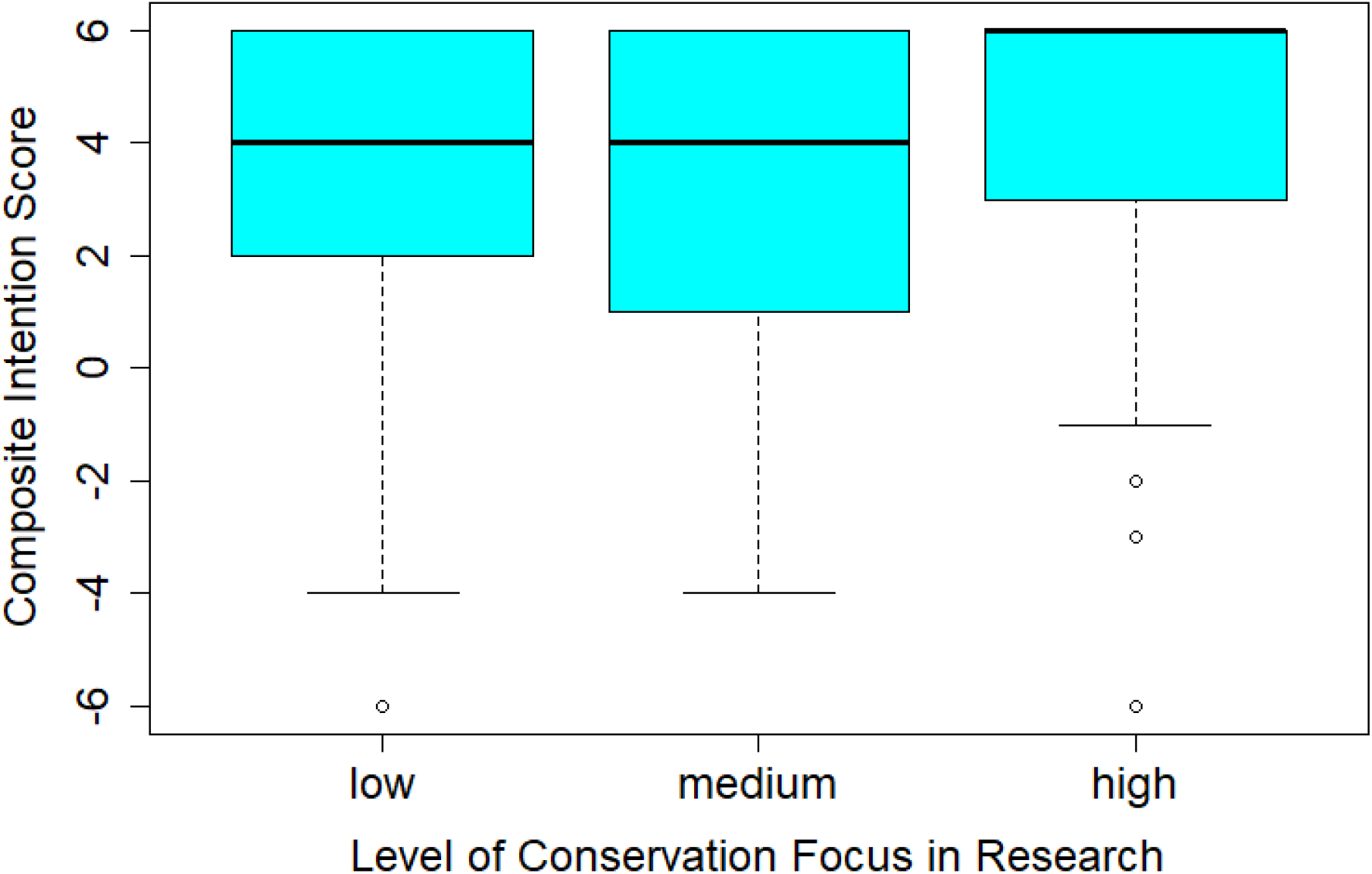
Boxplot of low (n=55), medium (n=23), and high (n=100) levels of conservation focus in sampled herpetologists’ research vs their intention to participate in public engagement for at least 10 hours over the next 12 months.

**Table 1.**
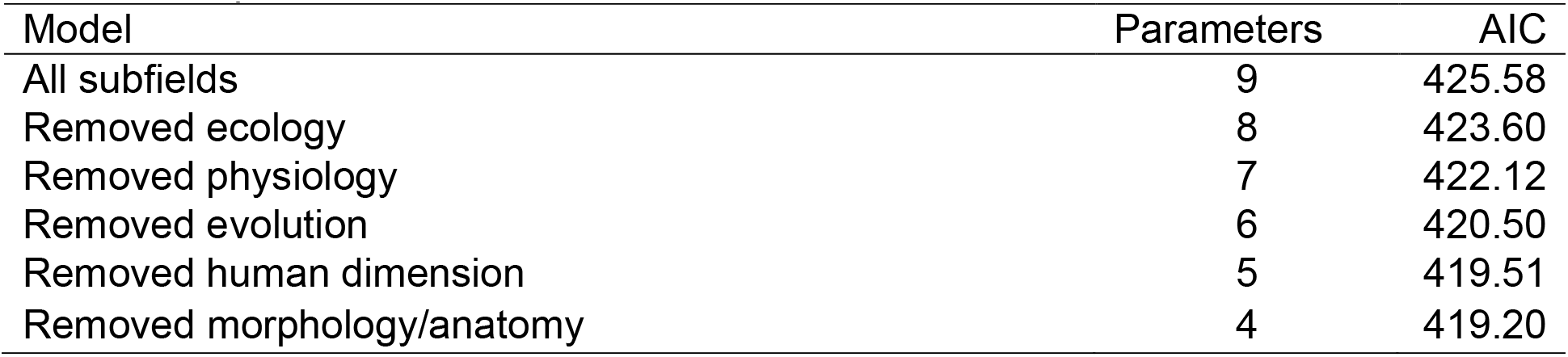
Stepwise model selection for

| Model | Parameters | AIC |
| --- | --- | --- |
| All subfields | 9 | 425.58 |
| Removed ecology | 8 | 423.60 |
| Removed physiology | 7 | 422.12 |
| Removed evolution | 6 | 420.50 |
| Removed human dimension | 5 | 419.51 |
| Removed morphology/anatomy | 4 | 419.20 |

**Table 2.** Final results for model parameters of ordinal logistic regression measuring hours of engagement of respondent herpetologists by subfield

|  | Value | Std. Error | t value | p value | Odds ratio | 95% CI |
| --- | --- | --- | --- | --- | --- | --- |
| Conservation Biology | 0.752 | 0.342 | 2.199 | 0.03 | 2.118 | 1.088 – 4.170 |
| Taxonomy | 0.833 | 0.359 | 2.319 | 0.02 | 0.939 | 1.430 – 4.687 |
| Behavior | 0.682 | 0.297 | 2.299 | 0.02 | 0.765 | 1.111 – 3.561 |
| Education | 1.114 | 0.569 | 1.958 | 0.05 | 1.309 | 0.988 – 9.348 |

## Discussion

Overall, we found that respondents were generally involved in public engagement activities, with less than 4% not participating in the past year. Similar patterns of general support and participation in public engagement activities have been noted in both general populations of scientists (Pew Research Center, 2015; Royal Society, 2006), as well as specific fields, such as particle physics (Rao, 2016); but not biology and physics professors at top research universities, where only 58% (n=150) of respondents were involved in science outreach (Ecklund et al., 2012). The Pew Research Center report (2015) also noted a similar pattern of engagement levels to our study, with the most responding scientists fitting in the “occasional participation” level (49%) as compared to our 52.2% participating in 10-49 hours in the past 12 months.

We also found that herpetologists were participating in some specific activities with potential for two-way communication at similar rates to scientists in other studies. For example, our study found that respondents used social media (56% vs 47%) and participated in policy making (20.7% vs 16%) at slightly higher rates than those in the AAAS study (Pew Research Center, 2015). Herpetologists in our study volunteered in schools (29.8%) more often than AAAS scientists (20%), but less often than biologists and physicists at top research universities (32%; Ecklund et al., 2012). Our respondents blogged less often than AAAS scientists (8.2% vs 24%). The small differences found between studies could be due to temporal changes such as an increase in general support for public engagement activities, federal funding for broader impacts, or a decrease in the popularity of written blogs since the 2015 AAAS study.

The connection between public engagement activities of scientists and public interest related to their field may help explain some of the patterns we found in this study, but additional factors related to wildlife outcomes may also encourage herpetologists to engage. Wildlife conservation is intimately tied to broader societal processes including socioeconomic factors, policy, and cultural norms; the field is directly tied to public interest. Therefore, we expected herpetologists who participated in conservation research to have higher levels of intention to participate in engagement activities. As we hypothesized, engagement intentions differed significantly in responding herpetologists based on their level of conservation focus in their own research. In addition, reporting conservation biology as one of their sub-fields was a significant predictor of how many hours herpetologists had engaged in public engagement in the last 12 months. While public interest is one possible explanation for the greater intentions seen among more conservation-oriented herpetologists, public opinions of reptile and amphibian conservation may also play a role. During qualitative interviews, we found that herpetologists’ interest in conserving herpetofauna is often tied to societal beliefs and the public’s negative attitudes toward and persecution of herpetofauna (Hecht, 2021). Thus at least for conservation-oriented herpetologists, scientific engagement activities appear to be driven in part by herpetofauna welfare rather than public interest. Thus, the influence of public interest on scientists’ engagement levels may expand beyond how much scientists feel the public is interested in their field to include how much the public impacts or is integrated with a scientist’s field of study.

Another finding of this study is that factors measuring the conservation focus of respondents’ research were not consistent in their predictive powers when comparing herpetologists’ future engagement intention to participation in the past 12 months. For example, a high level of conservation focus in research was associated with a higher intention to participate in engagement within the next 12 months, but did not result in higher levels of engagement within the past 12 months. In contrast, selection of conservation biology as a relevant sub-field of their research was a significant predictor of herpetologists’ engagement activity over the past 12 months, but not of their future intention to engage. In addition, taxonomy, behavior, and education were also significant predictors of future intenion. While this could suggest that barriers are preventing some individuals from participating at levels they would like to, alternative explanations are also likely. These sub-field findings may be due to survey design considerations for example, as respondents were able to report all sub-fields that they participate in, rather than only their main sub-field. Impacts of job sector may also be at play, as taxonomy is a common field of study associated with museums which often have a public mission focus.

While our survey respondents represented the overall demographics of herpetologists, ecology and conservation sub-fields may have been over-represented in our survey response. Since there are no data available about representation of sub-fields in herpetology, the implications of this to the results of our study are unclear. As those with a focus in conservation may be more likely to intend to participate or ultimately participate in public engagement based on the results of our study, one concern is that this finding suggests self-selection bias in our respondents toward those with increased interests in public engagement. Self-selection bias describes a sampling error where a certain sub-group of your population of interest, in this case those who may have different values or beliefs about public engagement, respond more often than those in other sub-groups, affecting survey results (Heckman, 1990; Whitehead, 1991). Despite these concerns, however, our respondents represented a normal distribution of actual participation in public engagement activities over the past 12 months. While this finding does not erase the possibility of self-selection bias based on interest, it does suggest that our results remain valid and still represent the spectrum our target population regarding their experiences with public engagement.

### Recommendations

We found that most respondents were already regularly engaging with the public with over 2/3 of respondents regularly engaging at least 10 hours of engagement activities over the last 12 months. Respondents also reported high levels of intention to participate in public engagement over the next 12 months, despite noted barriers. Public engagement seems to be generally accepted and valued by our respondents. However, the discrepancies noted between intention to engage and actual past activity as well as reports from participants that barriers prevented participation suggest that at least some herpetologists are interested in participating in public engagement activities more often than they are able. Therefore, we recommend that individuals and organizations with an interest in supporting public engagement activities focus on reducing the following key barriers and increasing awareness of engagement best practices to interested individuals.

Most of these respondents have not received training in public engagement skills or strategies, and many are interested in increasing their knowledge of best practices. Increasing herpetologists’ use of public engagement best practices may hold unique challenges since herpetologists may largely be unaware of the existence or importance of engagement best practices. Only a third of respondents received any type of training for conducting public engagement and most rarely or never consulted peer-reviewed literature or other resources to improve their engagement knowledge or skills. While these findings suggest that herpetologists may not be fluent in the language of public engagement, most respondents are doing something, suggesting that these respondents are not being prevented from engaging due to their lack of training or knowledge. However, the low use of some best practices, especially evaluation, shows that there is room for improvement in engagement practices, and it is unclear how effective their engagement practices may be. Therefore, one of the first challenges is to promote public engagement as a skill or sub-field much like other emerging conservation tools in herpetology such as structured decision making (Gregory et al., 2012).

Tempering these potential challenges in understanding and improving engagement best practices in herpetology, our respondents reported openness to learning more despite feeling it was unnecessary for their participation. Four out of five of herpetologist surveyed expressed interest in receiving some type of engagement training. Even more promising is that those responding expressed the most interest in receiving engagement training which also represented was one of the identified problem areas we found in our survey. Therefore, providing training opportunities in evaluation, developing engagement programs, and holding difficult conversations will likely receive interest, provided participants experience minimal time and money barriers to access these professional development opportunities. Providing these professional development opportunities at herpetology conferences or virtually may be one opportunity. Another option, due to the already strong culture of partnering for engagement opportunities, is to instead focus on connecting herpetologists to partners with expertise in these areas of interest to provide support.

Our last recommendations focus on reducing the barriers of time and money to allow herpetologists who wish to engage the opportunity to do so at their desired level. Herpetology and conservation organizations should consider developing funding opportunities designed to directly fund related engagement. Respondents were also interested in more engagement opportunities (23.0%), suggesting they may not be aware of already existing possibilities to participate. Conservation and herpetological organizations could focus on providing resources to connect herpetologists to public engagement opportunities and partnerships to increase the frequency and quality of engagement activities. Lastly, since most respondents were using their own resources to conduct engagement, employers interested in supporting their employee’s public engagement activities could allow some fraction of work time to be used for engagement activities or provide other resources.

Future research should further examine how much herpetologists are already utilizing best practices, especially for their desired goals. While our results show a few glaring issues with engagement best practices in herpetology, especially regarding evaluation, other best practices such as using live animals or adjusting messages to an audience were reported as being regularly utilized. However, since these questions were asked generally, and herpetologists may not share the same understanding of how to put these suggestions into practice, a more thorough investigation is necessary to know how to improve usage. For example, other studies have found that scientists frequently use a knowledge deficit model when communicating with the public (Davies, 2008), despite studies that suggest this model generally does not lead to positive outcomes in science communication or public engagement goals. If herpetologists are indeed adjusting their message based on their audience but only doing so in certain ways, like avoiding jargon, their approaches still largely focus only on transferring scientific knowledge to their audience, which may not necessarily lead to the positive outcomes best practices strive for.

While we recognize our results focus on herpetologists as a case study, we anticipate that these findings and recommendations may be more universally applicable in the conservation field, especially for individuals studying organisms or ecosystems that receive less conservation attention than charismatic animals, which generally receive more support for conservation both from the public and from within the conservation field itself (Clucas et al., 2008; de Pinho et al., 2014). Future research should investigate engagement in related sub-fields that have a natural resource and/or conservation focus to determine if our findings are unique to herpetology or are similar across related sub-fields. Specifically, these studies should compare sub-fields with more charismatic animals, like ornithology and mammalogy, with those sharing typically uncharismatic animals like entomology and shark biology. Comparisons of scientists working in basic ichthyology with those working in fisheries science, a sub-field with a long history of public involvement due to the recreational, economic, and subsistence nature of the field, could also provide additional insight on how public views of the study animal and the importance of the public in the field may influence scientists’ participation in public engagement.

## CRediT author statement

**K. Hecht:** Conceptualization, Methodology, Software, Formal Analysis, Investigation, Data Curation, Writing -Original Draft Preparation, Funding Acquisition **; K. Stofer:** Conceptualization, Methodology, Writing-Review & Editing, Supervision, Project Administration**; M. Monroe:** Conceptualization, Methodology, Writing-Review & Editing, Supervision**; G. Klarenberg:** Software, Formal Analysis, Writing-Review & Editing**; M. Nickerson:** Conceptualization, Writing-Review, Supervision.

